# Antibiotic tolerance, persistence, and resistance of the evolved minimal cell, *Mycoplasma mycoides* JCVI-Syn3B

**DOI:** 10.1101/2020.06.02.130641

**Authors:** Tahmina Hossain, Heather S. Deter, Eliza J. Peters, Nicholas C. Butzin

## Abstract

Antibiotic resistance is a growing problem, but bacteria can evade antibiotic treatment via tolerance and persistence. Antibiotic persisters are a small subpopulation of bacteria that tolerate antibiotics due to a physiologically dormant state. Hence, persistence is considered a major contributor to the evolution of antibiotic-resistant and relapsing infections. Here, we used the synthetically developed minimal cell *Mycoplasma mycoides* JCVI-Syn3B to examine essential mechanisms of antibiotic survival. The minimal cell contains only 473 genes, and most genes are essential. Its reduced complexity helps to reveal hidden phenomenon and fundamental biological principles can be explored because of less redundancy and feedback between systems compared to natural cells. We found that Syn3B evolves antibiotic resistance to different types of antibiotics expeditiously. The minimal cell also tolerates and persists against multiple antibiotics. It contains a few already identified persister-related genes, although lacking many systems previously linked to persistence (e.g. toxin-antitoxin systems, ribosome hibernation genes, etc.).

## Introduction

The evolution of antibiotic resistance is a pressing public health concern in the 21st century; antibiotic resistance results from one or more genetic mutations that counteract an antibiotic (Van den Bergh et al., 2016). Resistance is regarded as the primary culprit of antibiotic treatment failure, but bacteria also employ less publicized strategies to evade antibiotic treatment, namely antibiotic tolerance and persistence (Fisher et al., 2017; Harms et al., 2016; Lewis, 2010). Persisters are a subpopulation of tolerant cells, which can sustain longer against antibiotic treatment in comparison to slow-growing dying cells by entering a metabolically repressed state (non-multiplying cells) (Cabral et al., 2018; Fisher et al., 2017). Most antibiotics are only effective against growing cells, and by not growing, the persister population can survive longer even without being genetically resistant. Here, we define persisters based on a kill curve and the original paper where persistence was proposed (Bigger, 1944) (our definition of persistence is explained in detail in the discussion). Two types of persisters may exist, triggered persisters (formed by environmental stimuli) and spontaneous persisters (generated stochastically) (Balaban et al., 2019; Balaban et al., 2004; Sulaiman and Lam, 2020; Uruén et al., 2021), although spontaneous persister formation is controversial and evidence is sparse (Keren et al., 2004; Kim and Wood, 2016; Orman and Brynildsen, 2013). What makes persisters medically relevant is that they can revive and give rise to a new progeny after antibiotic treatment; the new progeny can be genetically identical to the original susceptible kin, and this process plays a pivotal role in recurring infections (Wilmaerts et al., 2019b) (Fig. 1). Furthermore, evolutionary studies have determined that repeated exposure to antibiotics over many generations rapidly increases tolerance leading to antibiotic resistance (Balaban and Liu, 2019; Cohen et al., 2013; Fridman et al., 2014; Liu et al., 2020; Schumacher et al., 2015; Sulaiman and Lam, 2020; Van den Bergh et al., 2016).

**Fig. 1.**
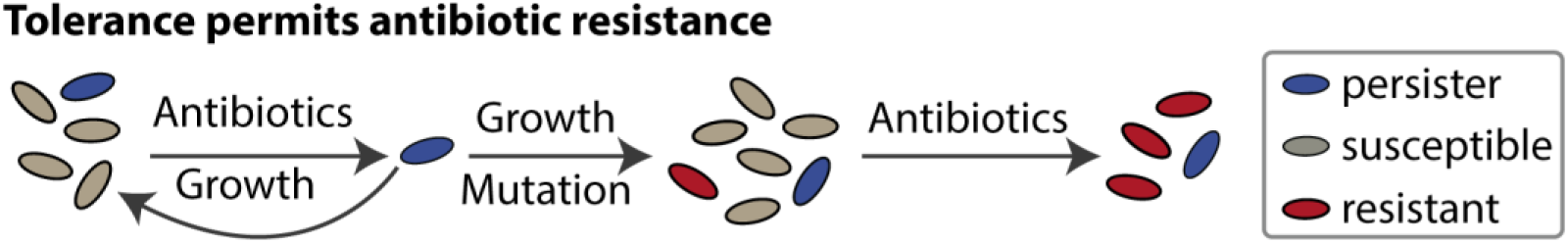
Persisters survive antibiotic treatment, reestablish the population when antibiotics are removed, and increase the odds of gaining resistance. Presumably, viable but nonculturable cells (VBNCs) can do this too (although we did not study VBNCs in this work).

The precise molecular mechanisms underlying persistence are still debated (multiple mechanisms are likely to exist), albeit several genes have been implicated (Wilmaerts et al., 2019a; Wu et al., 2015). Toxin-antitoxin (TA) systems have been implicated in persistence and were previously thought to be the key players for persistence (Lewis, 2010, 2012; Wang and Wood, 2011). Particularly important is the HipAB TA system, because a toxin mutant, *hipA7*, increases persistence by impeding cellular growth in the absence of its cognate antitoxin *hipB* (Moyed and Bertrand, 1983). Subsequent studies also report that overexpression of other TA systems’ toxins increased persistence (Correia et al., 2006; Kim and Wood, 2010; Korch and Hill, 2006). In contrast, new research found that the deletion of 10 TA systems (∼22% of TA system in the genome) in *E. coli* does not have an apparent effect on persistence both at the population and single-cell levels (Goormaghtigh et al., 2018), though *E. coli* can have more than 45 TA systems (Horesh et al., 2018; Karp et al., 2014; Xie et al., 2018). Though this work supports that TA systems are not controlling persistence, the remaining 35 TA systems (∼78%) could be controlling this phenomenon. Mauricio *et al*. studied persistence on Δ12TA *Salmonella enterica* and demonstrated that TA systems are dispensable (Pontes and Groisman, 2019), although this strain has 18 predicted TA systems based on the TAfinder tool (Xie et al., 2018). Another knockout study on *Staphylococcus aureus* deleted three copies of type II TA system from the Newman strain (Conlon et al., 2016). But further studies identified several type-I (sprG1/sprF1, sprG2/sprF2, sprG4/sprF4) (Riffaud et al., 2019) and type-II (SavRS) (Wen et al., 2018) TA system in HG003 and NCTC-8325 strains, respectively. These are the parental strains of Newman, and these TA genes are also found in the Newman strain (Sassi et al., 2015). Moreover, a recent study showed that antitoxin sprF1 mediates translation inhibition to promote persister formation in *S. aureus* (Pinel-Marie et al., 2021). These findings raise many questions about TA systems’ implication in bacterial persistence (Goormaghtigh et al., 2018; Kim and Wood, 2016; Pontes and Groisman, 2019; Tsilibaris et al., 2007). There are major limitations of studying TA systems in native bacteria due to their high abundance in most bacterial genomes; they often have redundant and overlapping functions and can have interdependencies within network clusters (Harms et al., 2018; Ronneau and Helaine, 2019; Wang et al., 2013).

Additionally, TA systems respond to a variety of stresses (e.g. bacteriophage infection, oxidative stress, etc.), which creates a hurdle to probe their connection with any phenomenon related to stress response, namely antibiotic tolerance and persistence (Harms et al., 2018; Kang et al., 2018; Ronneau and Helaine, 2019). TA systems are also involved in other mechanisms in the cell that respond to stress, such as the stringent and SOS responses (Ronneau and Helaine, 2019), virulence (De la Cruz et al., 2013), and the regulation of pathogen intracellular lifestyle in varied host cell types (Lobato-Marquez et al., 2015). Furthermore, new types of TA systems may be yet unidentified. These challenges could be resolved using a strain that lacks canonical TA systems. But TA systems are naturally abundant, and large-scale knockouts are both error-prone and labor-intensive. We took advantage of the recently developed minimal cell that does not encode any sequences displaying homology to known TA systems and showed that it can still form persisters.

Several other mechanisms have also been considered in persister research including SOS response, oxidative stress response, etc. (Trastoy et al., 2018; Wilmaerts et al., 2019a; Wilmaerts et al., 2019b). Two extensively studied phenomenon related to persistence are cellular ATP levels and (p)ppGpp levels. The accumulation of (p)ppGpp (stress sensing alarmone) mediates the stringent response, which controls a stress-related persistence mechanism (Harms et al., 2016). (p)ppGpp regulates many networks (such as ribosome dimerization) that can cause cells to go into dormancy (Gaca et al., 2015; Song and Wood, 2020; Wood and Song, 2020). Another well-studied model, *ATP depletion increases persistence*, has drawn much attention in persister research (Conlon et al., 2016). This finding is consistent with REF (Pu et al., 2019), which demonstrated that lower ATP levels lead to protein aggregation and increased tolerance. This result is also coherent with our recently published data, which established that interfering with protein degradation by forming a proteolytic queue at ClpXP will increase tolerance levels dramatically (Deter et al., 2019a). Transcriptomic analysis of queuing-tolerant population showed upregulation of genes related to metabolism and energy (Deter et al., 2020b). However, other studies reported that ppGpp and ATP reduction are not essential for persistence (Bhaskar et al., 2018; Chowdhury et al., 2016; Pontes and Groisman, 2019). These contradictory studies are common in persister research, and we hypothesize these inconsistencies are due to the interconnection of gene networks surrounding persistence.

One goal of our study is to clarify mechanisms using a minimal, simpler system. Research over the last several years has resulted in a lot of discussion concerning genes essential for persistence (Pontes and Groisman, 2019; Ronneau and Helaine, 2019; Wilmaerts et al., 2019a; Wilmaerts et al., 2019b). Since persistence and tolerance are present in phylogenetically diversified bacterial species (Meylan et al., 2018; Wilmaerts et al., 2019b), it is feasible that an underlying genetic mechanism is conserved in evolutionarily related microorganisms, and the most likely candidates are essential genes or the disruption of crucial networks. In our recent work, we demonstrate that antibiotic tolerance in *Escherichia coli* may result from a whole-cell response and can occur through multiple pathways or networks, which may work simultaneously and cooperatively to survive against antibiotics (Deter et al., 2020b).

Our strategy to study the underlying mechanisms of persistence was to use a bacterial species that contains mainly essential genes and networks with reduced complexity and fewer networks. The minimal system can reveal hidden phenomena (Glass et al., 2017) of antibiotic survival. For example, genes and networks that have previously been identified can be eliminated as causal if the genome lacks them. In contrast, genes present in Syn3B that were previously identified in other organisms can become the focus of the work. The reduced complexity has its limits as there are likely several methods microbes use to survive antibiotics, and not all methods will be in a minimal system. With these limitations well-understood, we explored antibiotic tolerance, persistence, and resistance in the minimal cell *Mycoplasma mycoides* JCVI-Syn3B (called Syn3B throughout), a synthetic genetic background that contains the least number of genes and smallest genome of any known free-living organism (Syn3B contains 473 genes and its genome is ∼531 Kbp long, while *E. coli* contains >4,000 genes and its genome is ∼4,600 Kbp long). The Syn3B genome was minimized from Syn1.0 (contains a chemically synthesized genome of *M. mycoides* subspecies capri with some watermarks and vector sequences) by removing non-essential genes (Gibson et al., 2010; Hutchison et al., 2016). Syn3B consists predominantly of essential and a few non-essential (added for ease of genome manipulation) and quasi-essential genes (required for robust growth) (Hutchison et al., 2016). This microbe was designed to have a minimal number of genes and networks with the expectation that many of the first principles of cellular life could be explored in the simplest biological systems (Glass et al., 2017). We show that Syn3B populations contain both persister and tolerant cells, and its genome contains a few previously identified persister-related genes. This work establishes that many systems previously shown to be related to bacterial persistence, such as TA systems and ribosome dimerization, are not essential for persistence in the minimal cell, and it demonstrates the effectiveness of using the minimal cell to study antibiotic survival.

## Result

### Whole-genome analysis of the evolved minimal cell

Although the minimal genome (Syn3B) was designed for ease of genetic manipulation, Syn3B grows slowly and to a low cell density. We adjusted the original SP4 media (Tully et al., 1979) to address these limitations. The new media, SP16, allows for faster cell growth and higher yields in liquid cultures. We noticed that Syn3B growth was slightly better after every subculture. Thus, we did a cyclic subculture of Syn3B in our optimized media by passing cells during logarithmic growth. After 26 passages, we observed better growth and isolated a single colony named Syn3B P026 (<u>P</u>ass <u>26</u>). This strain has a shorter lag phase and an increased growth rate; the average doubling time of P026 is approximately 2.5 hours compared to the ancestral Syn3B doubling time of ∼6 hours under the same conditions (Fig. S1.A-B). We performed a whole-genome analysis of Syn3B P026 to examine the genetic basis of these changes. Most of the mutations (9 of 11) in P026 are intergenic except one synonymous (*fakA*) and one non-synonymous (*dnaA*) (Table 1). *fakA* is a fatty acid kinase, and there is less evidence to suggest a direct connection to substantial alterations in bacterial growth. *dnaA* is a positive regulator of DNA replication initiation. Considering that bacterial growth rate is dependent on the frequency of DNA replication and that *dnaA* mutants have been known to result in over-replication (Skarstad and Boye, 1994), we hypothesize that the mutation of *dnaA* in P026 could be responsible for the higher growth rate. We also sequenced the *dnaA* gene from 5 colonies of a P026 subculture, and they had similar growth rates as the P026 culture we initially sequenced. All 6 colonies have the same mutation in the *dnaA* gene. These results suggest that the *dnaA* is the likely cause for the increased growth rate of the evolved strain P026 compared to the parent strain, but we did not further pursue the role of *dnaA* because it is not the main focus of this study.

**Table 1.**
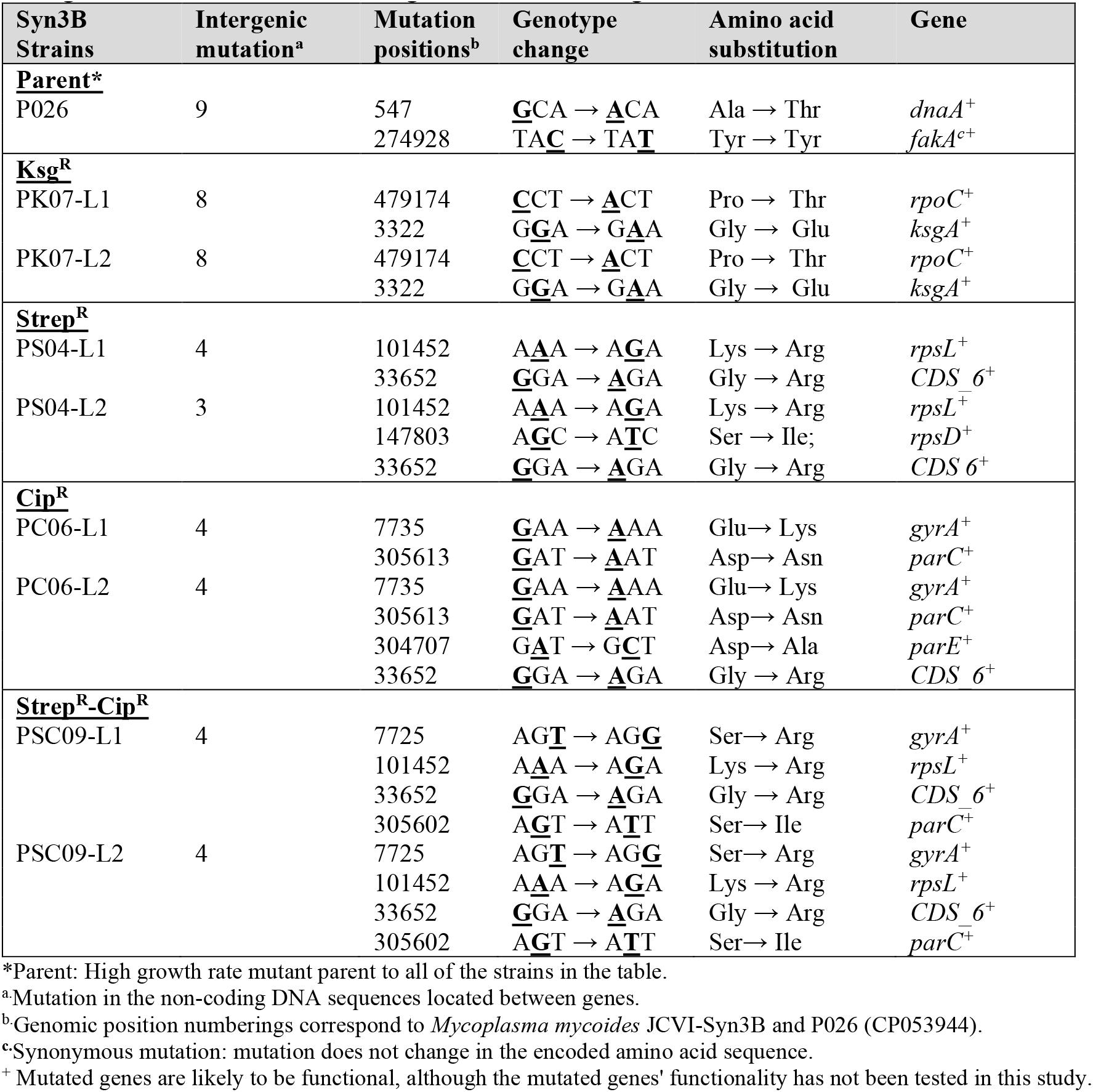
Mutation of *Mycoplasma mycoides* JCVI-Syn3B, and Whole Genome Sequence (WGS) analysis identified the mutations (bold and underlined). All antibiotic-resistant mutant strains were named based on the number of passes (P) in a specific antibiotic (K: Ksg, S: Strep, C: Cip, SC: Strep-Cip). All antibiotic-resistant mutants were selected from two separate evolutionary lineages, and named L1 for lineage 1 or L2 for linage 2.

### Syn3B and evolved minimal cell P026 display antibiotic tolerance and persistence

We assessed tolerance and persistence (Fig. 2) of Syn3B (parent strain) and Syn3B P026 cultures using two bactericidal antibiotics, ciprofloxacin (a fluoroquinolone) and streptomycin (an aminoglycoside). Ciprofloxacin inhibits DNA replication by targeting DNA gyrase and topoisomerase IV activity (Sanders, 1988). Streptomycin blocks protein synthesis by irreversibly binding to the 16s rRNA of the 30S ribosomal subunit (Luzzatto et al., 1968). Syn3B and P026 were grown to stationary phase and diluted into fresh media containing the antibiotics to observe population decay (see Methods). Both parent strain and P026 cultures showed a typical biphasic death curve from stationary phase; the death rate became slower at ∼20-24 h treatment with both antibiotics compared to the earlier stage of population decay, which indicates Syn3B displays persistence (Fig. 2. B-C). It appears that the survival was higher in the Syn3B ancestor strain than P026 for both antibiotic treatments. We posit that it could be due to the slower growth rate of ancestor strain, which is consistent with the literature, as several research groups already establish that growth rate has a strong correlation with antibiotic susceptibility (Abshire and Neidhardt, 1993; Lee et al., 2018; Pontes and Groisman, 2019; Tuomanen et al., 1986). As we observed tolerance and persistence in both the ancestor strain and Syn3B P026, we decided to do further analysis with Syn3B P026, because it was much easier to work.

**Fig. 2.**
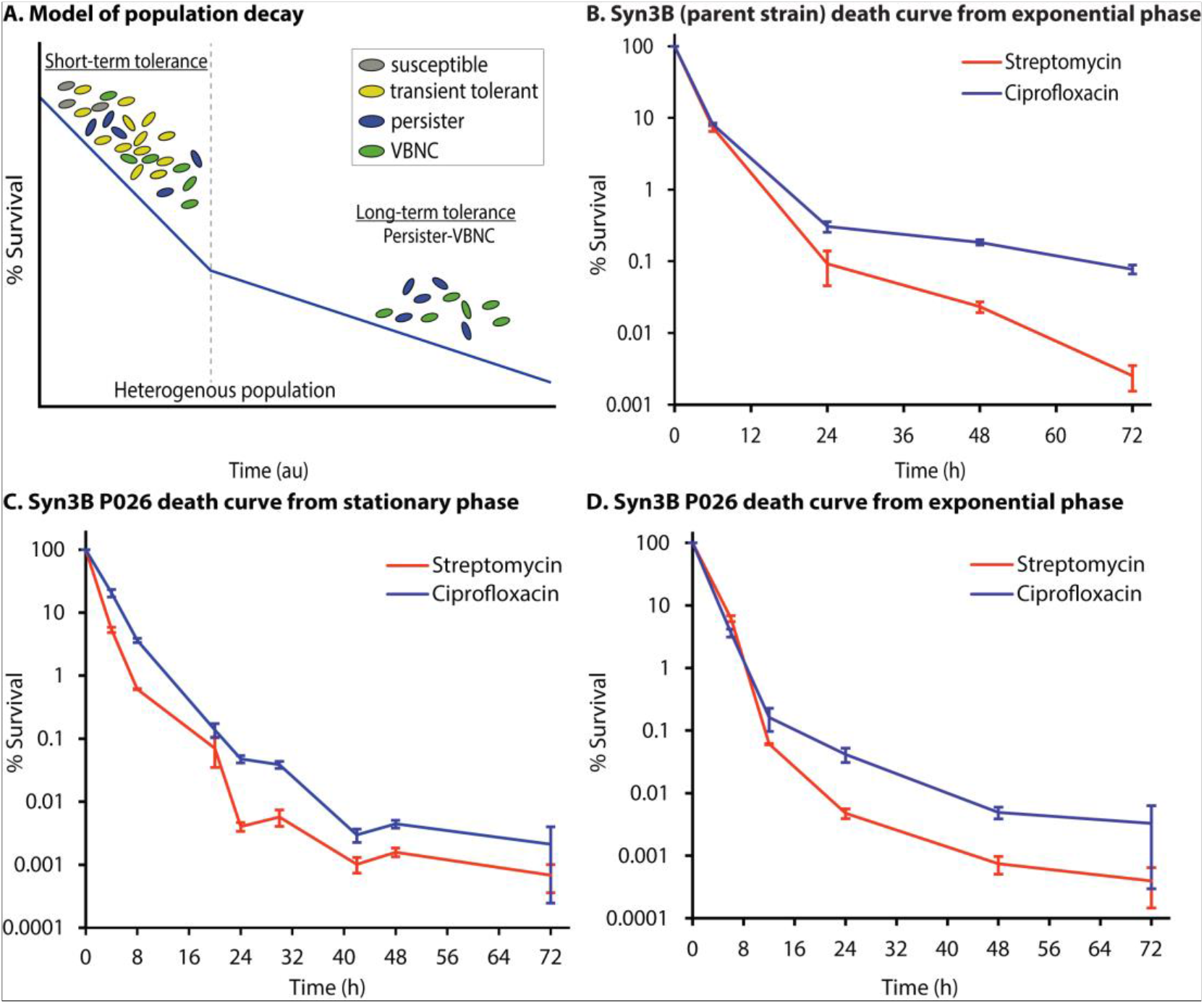
Population decay during antibiotic treatment shows Syn3B persistence. **A**. A simplified model of population decay having two phases: short-term tolerance phase and long-term tolerance phase. Both phases contain heterogeneous populations. The short-term tolerant phase contains susceptible cells, slow-growing cells, transient tolerant cells, persister and viable but nonculturable cells (VBNCs). Transient tolerant cells survive longer than fast-growing susceptible cells, while transient tolerant cells die quicker than the persisters and VBNCs (VBNCs and persisters were not distinguished in this study). The long-term tolerant phase contains both persisters and VBNCs. **B**. Population decay of Syn3B (parent strain). Overnight cultures of Syn3B (parent strain) were grown to stationary phase, diluted to 1:10, and then treated with streptomycin (100 µg/mL) or ciprofloxacin (1 µg/mL) and sampled over time. Error bars represent SEM (n ≥ 3). **C-D**. Population decay of Syn3B P026. Syn3B P026 cells were treated with streptomycin (100 µg/mL) or ciprofloxacin (1 µg/mL) and sampled over time. (**C)** Stationary phase cells were diluted 1/10 into fresh media containing antibiotics. Error bars represent SEM (n ≥ 6). **(D)** Exponential phase cells were treated with streptomycin (100 µg/mL) or ciprofloxacin (1 µg/mL) and sampled over time. Error bars represent SEM (n ≥ 3). There is 100% survival at time zero because percent survival is determined by the surviving CFU/ml compared to the CFU/ml at time zero.

Persister cells have been identified in native bacterial species in both stationary and exponential phase cultures, and we tested if this was also true for the minimal cell. P026 was grown to exponential phase and treated with antibiotics to determine whether the minimal genome showed a similar biphasic death curve in exponential phase. As expected, we observed a similar biphasic killing curve in exponential phase with slightly lower survival in exponential phase compared to stationary phase for both antibiotics (Fig. 2. D). Moreover, we performed resistance assays through the course of this work to rule out the possibility of resistance instead of tolerance or persistence (see Methods), and no resistant colonies were identified. We then tested if the surviving population had increased persister levels after antibiotic treatment. After 48 h of antibiotic treatment, the culture was passed into fresh media and then grown to stationary phase. The culture was then exposed to the same antibiotic under the same condition, and as expected, no significant difference between the two death curves was observed (Fig. S2).

At this point, we have demonstrated that tolerance and persistence can be detected in Syn3B cultures during both stationary and exponential phase, and this is consistent with native bacterial species. Another well-known phenotype of antibiotic survival is that the tolerant population increases with co-treatment of bacteriostatic and bactericidal antibiotics (Lewin and Smith, 1988) (pre-treatment is another method (Kwan et al., 2013)). To use the minimal cell as a model of antibiotic survival, this phenomenon should also be observed. Bacteriostatic antibiotics can counteract the bactericidal antibiotic’s killing activity by arresting the growth of rapidly growing cells, which increases tolerance to the antibiotics (Kwan et al., 2013). Chloramphenicol is a bacteriostatic antibiotic that inhibits translation by binding to bacterial ribosomes and inhibiting protein synthesis, thereby inhibiting bacterial growth (Volkov et al., 2019). We co-treated with the bacteriostatic antibiotic chloramphenicol and either streptomycin or ciprofloxacin. As expected, the overall percent survival with chloramphenicol co-treatment increased compared to streptomycin or ciprofloxacin alone after 24 h and 48 h (Fig. 3). These results support that inhibition of translation by co-treating the cell with chloramphenicol increases tolerance in P026.

**Fig. 3.**
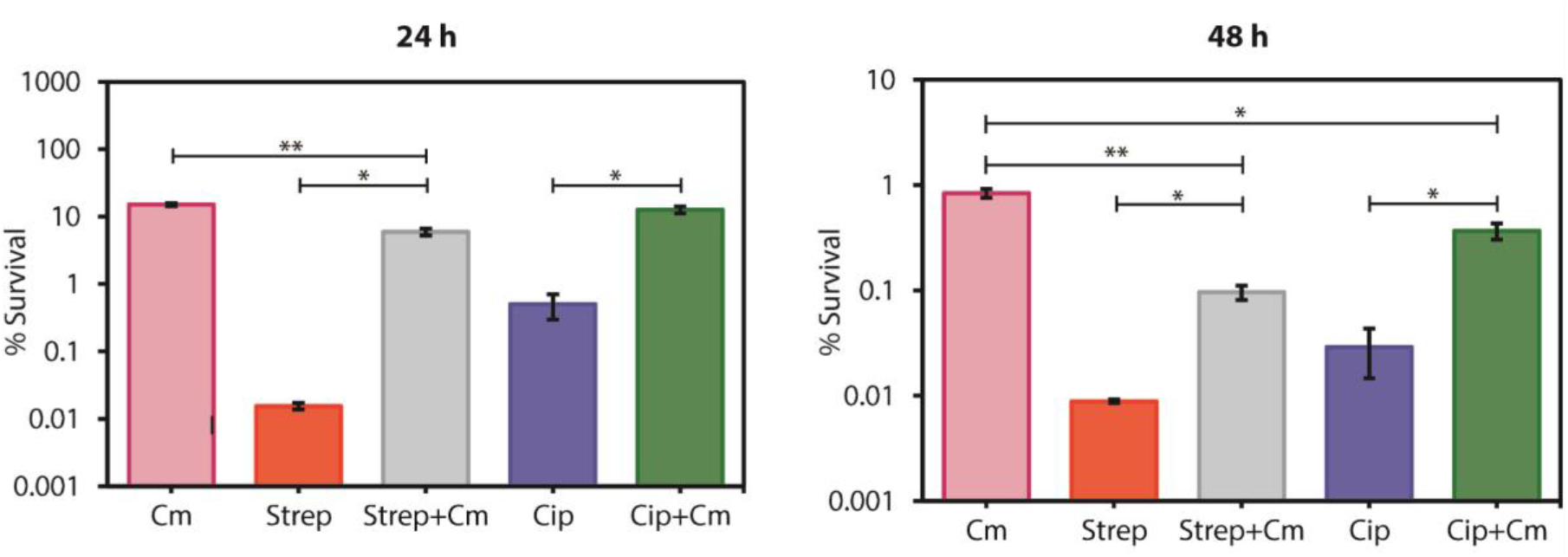
The co-treatment of bactericidal antibiotic (streptomycin (Strep) or ciprofloxacin (Cip) with a bacteriostatic antibiotic (chloramphenicol (Cm)) shows increased survival of cells for 24 h (left) and 48 h (right) in Syn3B P026. Error bars represent SEM (n ≥ 3). *p<0.05. **p<0.01.

### The minimal genome contains previously identified genes related to tolerance and persistence

The mechanisms of persister formation are complex, but recent studies identify several genes involved in this process (Cameron et al., 2018; Wilmaerts et al., 2019a). A few of these genes are present in the minimal genome. We identified known and predicted genes related to tolerance and persistence in the Syn3B genome (Table 2). During nutrient limitation, bacteria generally switch their gene expression profile from rapid growth to a survival state by (p)ppGpp levels (a hallmark of the stringent response), which is regulated by the RelA protein in *E. coli* (Boutte and Crosson, 2013). Another important gene in the stringent response is *phoU*, a global negative regulator, which is deactivated upon inorganic phosphate (Pi) starvation. Mutation or deletion of this gene in other microorganisms leads to a metabolically hyperactive state and decreased persistence (Li and Zhang, 2007; Zhang and Li, 2010). The SOS response is another signaling pathway that upregulates DNA repair functions, which appears to be linked to bacterial persistence. Genes related to the SOS response have been found upregulated in *E. coli* persister (e.g. *uvrA, uvrB*) (Keren et al., 2004), and other mutants (such as *xseB*) have lost their ability to induce detectable levels of persistence when pretreated with rifampicin, a bacteriostatic antibiotic (Cui et al., 2018).

**Table 2.** Annotation of mutated genes from Table 1.

| Mutated Gene | Annotation |  |
| --- | --- | --- |
| *dnaA*<sup>+</sup> | Chromosomal replication initiator protein DnaA | 201 |
| *fakA*<sup>+</sup> | Dihydroxyacetone kinase | 202 |
| *rpoC*<sup>+</sup> | DNA-directed RNA polymerase subunit beta | 203 |
| *ksgA*<sup>+</sup> | 16S rRNA (adenine(1518)-N(6)/adenine(1519)-N(6))- dimethyltransferase | 204 |
| *rpsL*<sup>+</sup> | 30S ribosomal protein S12 | 205 |
| *CDS_6*<sup>+</sup> | Unknown | 206 |
| *rpsD*<sup>+</sup> | 30S ribosomal protein S4 | 207 |
| *gyrA*<sup>+</sup> | DNA gyrase subunit A | 208 |
| *parC*<sup>+</sup> | DNA topoisomerase 4 subunit A | 209 |
| *parE*<sup>+</sup> | DNA topoisomerase 4 subunit B | 210 |

Other literature advocates the connection of persistence with energy metabolism, although the outcomes are often inconsistent between models (Wilmaerts et al., 2019a). For example, the deletion of *atpA* (encoding for ATP synthase subunit alpha) decreases the persister fraction (Lobritz et al., 2015), but deletion of genes encoding other parts of the ATP synthase (*atpC and atpF*) increases the persister fraction (Girgis et al., 2012; Kiss, 2000). Several heat shock proteins, mainly proteases (e.g. *lon* (Pu et al., 2016; Wu et al., 2015)) and chaperons (e.g. *dnaK* (Wu et al., 2015), *dnaJ* (Hansen et al., 2008), *clpB* (Wu et al., 2015)), are related to persistence considering that deletion of those genes decreases persistence.

Another survival mechanism connected to persistence is trans-translation, which allows bacteria to recycle stalled ribosomes and tag unfinished polypeptides for degradation, which in *E. coli* requires tmRNA (encoded by *ssrA*) and a small protein (smpB). The deletion of these genes causes persister-reduction in *E*.*coli* (Liu et al., 2017; Wu et al., 2015; Yamasaki et al., 2020). Additional genes (*glyA* (Cui et al., 2018), *galU* (Girgis et al., 2012), *trmE* (Cui et al., 2018), *efp* (Cui et al., 2018), *ybeY* (Cui et al., 2018), etc.; see Table 2) are indirectly related to stress response or metabolism and have been reported to affect persistence. Overall, this shortlist (Table 2 compare to Table S1) of genes demonstrates that there are far fewer genes to explore in the minimal cell than any other known free-living microorganism on the planet. However, further exploration is needed to test if there is a relationship between genes in Table 2 to antibiotic survival in the minimal cell.

### Evolution of the minimal cell against different types of antibiotic and whole-genome analysis of the resistant strain

Up until this point, we have focused on antibiotic tolerance and persistence. For the minimal system to be also useful as a model of antibiotic resistance, it must be able to evolve resistance without horizontal gene transfer (i.e. isolated from other organisms). We hypothesize that due to lack of complexity and less control of mutation, simpler cells likely evolve faster than more complex organisms; a simpler, slimmed-down genome allows for rapid evolution to selective pressures. We selected different classes of bactericidal antibiotics including streptomycin (Strep), ciprofloxacin (Cip), a combination of streptomycin and ciprofloxacin (Strep-Cip), and one bacteriostatic antibiotic-kasugamycin (Ksg), to study the evolution of antibiotic resistance in the minimal genome. We applied cyclic antibiotic treatments where we repeatedly regrew and retreated the culture with the lethal dose of the same antibiotic for a few cycles with two biological replicates (Fig. S3B.) to make resistant mutants. In seven passes or less (nine passes with two antibiotics), the minimal genome rapidly evolved resistance to all antibiotics tested. We isolated single colonies (two separate evolutionary lineages named L1 for lineage 1 or L2 for linage 2) from each resistant mutants named Syn3B PS04 (<u>4</u> Passes in <u>S</u>treptomycin), PC06 (<u>6</u> Passes in <u>C</u>iprofloxacin), PSC09 (<u>9</u> Passes in <u>S</u>treptomycin and <u>C</u>iprofloxacin), and PK07 (<u>7</u> Passes in <u>K</u>asugamycin). We then analyzed the whole genome for mutations. Most mutations were in intergenic regions of the resistant mutants, similar to Syn3B P026 (Table 1 and 2).

We observed Syn3B P026 quickly evolved against streptomycin, as it only took 4 passes to become resistant. Only two non-synonymous mutations were found in both lineages. The first one is in the *rpsL* gene (encoding the S12 ribosomal protein), which is frequently identified in streptomycin-resistant strains (Villellas et al., 2013), and the other one is the *CDS_6* gene, the function of which is not yet known. In PS04_L2, we also observed another mutation in the *rpsD* gene (encoding the S4 ribosomal protein). We found that Syn3B P026 took 6 and 9 cycles to gain resistance against ciprofloxacin and a combination of both streptomycin and ciprofloxacin, respectively. In both replicates of PC06, we observed mutations in *gyrA* (encoding the DNA gyrase subunit A) and *parC* (encoding DNA topoisomerase 4 subunit A). In PC06_L2, we observed another two mutations, one in *parE* (encoding DNA topoisomerase 4 subunit B) and another in the *CDS_6* gene. For the combined streptomycin-ciprofloxacin resistant strains, both lineages showed similar mutations in *rpsL, gyrA, parC*, and *CDS_6* genes. In the kasugamycin-resistant strain PK07, only two nonsynonymous mutations were found. The first one is in the *ksgA* gene, which encodes a methylase that modifies 16S rRNA, and inhibition of this protein by the antibiotic kasugamycin causes susceptibility. Therefore, when *ksgA* is inactivated, cells became kasugamycin-resistant (Mariscal et al., 2018; van Gemen et al., 1987). We reasoned that the nonsynonymous mutation in *ksgA* in PK07 makes the protein inactive and able to confer kasugamycin resistance. The other detected non-synonymous mutation is in *rpoC* (RNA polymerase subunit).

## Discussion

Overall, we have demonstrated that the minimal cell contains both antibiotic tolerant and persistent subpopulations, and it can quickly evolve to gain resistance. We have successfully generated an evolved strain of the minimal cell Syn3B P026, which has a better growth rate than the ancestral strain and overcomes one of the major difficulties of working with the minimal cell. We show that Syn3B can be used as a model for studying antibiotic survival, especially when we consider that the minimal genome’s core proteins are present in many other microorganisms including human pathogens. Syn3B has over 50% similarity to human pathogens that have been identified as a concern with respect to the growing antibiotic-resistance problem (Table S2) (Centers for Disease Control and Preventions, 2019). For example, out of 455 protein-encoding genes of Syn3B, 338 of them (74%) have homologs to *S. aureus* NCTC 8325 proteins. *S. aureus* causes meningitis and pneumonia, and antibiotic resistance is a major problem of this organism (Foster, 2017).

The definition of persistence is often debated, but we argue that the original definition proposed by Biggers in his seminal 1944 paper (Bigger, 1944), where persisters were first classified and named, is sufficient. Syn3B meets 9 out of 10 characteristics defined by Bigger. The only characteristic not found in Syn3B (point 4 in Table 4) was an untested hypothesis that Bigger put forward about persisters, which states that persisters can be induced without antibiotic stress (now known as spontaneous persisters). This characteristic has yet to be resolved and was not addressed in our work. Bigger made it clear that persister identification should occur over several days and suggested 72 h (Bigger, 1944). We tested Syn3B for 72 h (Fig. 2B-D), and it formed persisters. Though we agree with Bigger’s original assertion that prolonged antibiotic treatment is required to demonstrate persistence, the originally 72 h cutoff does not consider different dividing times of bacteria, which is likely to affect killing rates.

**Table 3.** Syn3B contains genes previously shown to be related to *E*.*coli* persistence and tolerance.

| Gene | Protein/RNA | Major function | REF |
| --- | --- | --- | --- |
| *dnaK, dnaJ, clpB* | Chaperon | Global regulator | (Hansen et al., 2008; Wu et al., 2015) |
| *phoU* | Phosphate-specific transport system accessory protein | Global regulator | (Li and Zhang, 2007) |
| *xseB* | Exodeoxyribonuclease | DNA mismatch repair and recombination | (Cui et al., 2018) |
| *uvrA, uvrB* | Endonuclease | SOS response | (Cui et al., 2018; Keren et al., 2004; Wu et al., 2015) |
| *relA* | GTP pyrophosphokinase | Stringent response | (Wu et al., 2015) |
| *lon* | Protease | Protease | (Wu et al., 2015) |
| *glyA* | Serine hydroxymethyltransferase | Metabolism | (Cui et al., 2018) |
| *atpA, atpC, atpF* | ATP synthase | ATP synthesis | (Girgis et al., 2012; Kiss, 2000; Lobritz et al., 2015) |
| *galU* | UTP--glucose-1-phosphate uridylyltransferase | Lipopolysaccharide precursor synthesis | (Girgis et al., 2012) |
| *trmE; efp; ybeY* | tRNA modification GTPase; Elongation factor P; Endoribonuclease | Translation | (Cui et al., 2018) |
| *ssrA; smpB* | Transfer messenger RNA (tmRNA); Small Protein B | Trans-translation | (Wu et al., 2015; Yamasaki et al., 2020) |

**Table 4.** In Bigger’s 1944 paper, he identified cells that survived antibiotics longer, named them persisters, and listed 10 characteristics of the persister subpopulation (Bigger, 1944). Syn3B meets 9 out of 10 characteristics defined by Bigger. Left column: summary of Bigger’s definition. Right column: similar characteristics of Syn3B persisters. We provided no evidence related to point 4 but observed a similar phenotype as Bigger to support point 10.

|  | **Bigger, 1944 persister definition** | **Syn3B persisters** |
| --- | --- | --- |
| 1. | An antibiotic failed to kill a small population of bacteria. | Multiple antibiotics failed to kill a small population of Syn3B. |
| 2. | Persisters are a small population initially present in the population. | Syn3B contains a small persister subpopulation among the susceptible population. |
| 3. | Persister can be induced or changed based on the environment. | The persistence (surviving population) level is different when the culture originated from a stationary phase or exponential phase. The persistence level is different when cultures were co-treated with bacteriostatic and bactericidal antibiotics compared to the bactericidal antibiotic only; chloramphenicol and streptomycin, and chloramphenicol and ciprofloxacin, compared to the antibiotics alone. |
| 4.* | *Untested hypothesis:* No evidence showed that persisters were produced by the presence of an antibiotic. | This was not tested for Syn3B persisters. |
| 5. | Bacteriocidal concentrations of an antibiotic killed the susceptible population, but not persisters. | The Syn3B susceptible population was killed by bacteriocidal concentrations of more than one antibiotic type, but persisters were not. |
| 6. | Persisters are likely insensitive to an antibiotic because they are dormant and non-dividing. The antibiotic used, penicillin, kills bacteria only when they divide. | Syn3B persisters also appear to be dormant and likely non-dividing, because the antibiotics used are more effective against dividing cells. |
| 7. | Descendants of persisters are no more resistant to an antibiotic than the original population. | Syn3B descendants of persisters are no more resistant to an antibiotic than the original population. |
| 8. | When the antibiotic is destroyed, the majority of persisters will emerge and grow normally. | Syn3B persisters grew once the antibiotic was removed or diluted below the MIC. |
| 9. | The antibiotic prolongs the dormant phase but not indefinitely. | Syn3B persisters continue to die, but slowly, the longer they are exposed to antibiotics. There is no plateau but a slow decline in cell death. |
| 10.* | *Untested hypothesis:* Persisters may be coming out of the dormant state, attempting to divide and being killed by the antibiotic. | This was not tested for Syn3B persisters. Syn3B persisters continue to die, but slowly, the longer they are exposed to antibiotics. There is no plateau but a slow decline in cell death. |

\* Points 4 and 10 were untested hypotheses of Bigger and were not tested in our work.

A more recent definition of tolerance and persistence was put forth by a consensus statement released after a discussion panel with 121 researchers (Balaban et al., 2019). They defined persistence similarly to the original Bigger’s paper with some slight changes. They defined persistence as a subpopulation of tolerance, and this is generally agreed upon in the literature. They then stated that the “tolerance state” could be distinguished from the “persistence state” based on a killing curve. We agree that persisters can be distinguished from other populations using a kill curve (Fig. 2A), the Bigger paper also demonstrated the importance of prolonged antibiotic treatment to identify persisters (Bigger, 1944), and we showed that Syn3B also has a distinguished death curve (Fig. 2B-D). However, we do not call the first death phase “tolerance,” as proposed by REF (Balaban et al., 2019), because naming this subpopulation tolerance is easily confused when the entire population is tolerant. We, instead, label the first phase (faster death rate) of the death curve as short-term tolerance phase contains heterogeneous population includes susceptible cells, transient tolerant cells (caused by slow-growth or heterogeneity in gene expression (El Meouche and Dunlop, 2018; Lee et al., 2019)), persisters and VBNCs. Susceptible cells are included because it will take time to kill off these cells regardless of the antibiotic used. Transient tolerant cells can tolerate antibiotics longer than susceptible cells, but not as long as VBNCs or persisters (VBNCs were not investigated in this study). We termed the second phase (slower death rate) as a long-term tolerance phase containing persister and VBNCs. We described in our definition that a death curve can distinguish short-term tolerance and persistence.

Few researchers have put forth that the definition for persisters should include a rapid decline in the short-term tolerance stage and then a persister plateau (Bartell et al., 2020; Mulcahy et al., 2010; Sahukhal et al., 2017; Song and Wood, 2021). Depiction of persisters as a plateau may underrepresent the heterogeneity of the population (Kaldalu et al., 2016), and a true plateau have a slope of zero for an extended period of time, which we do not see in our Syn3B and *E*.*coli* data. We first tested if a plateau is present in the model organism *E. coli*. Since other researchers (Aedo et al., 2019; Mohiuddin et al., 2020; Song et al., 2019) and ourselves (Deter et al., 2020a) have shown that *E. coli* forms persisters after three hours of ampicillin treatment at bactericidal concentrations (100 µg/mL treatment), we reanalyzed our previously published data (Deter et al., 2020a) and included 24 ampicillin exposure data to check for a plateau using a simple t-test point comparison. The hypothesis is there is no plateau, the slope of the line is not zero, and a slow population decay after 3 h. The null hypothesis is there is a plateau, the slope of the line is zero, and the population reaches a steady-state with no decay after 3 h. Comparing the p-values of 3 h ampicillin exposure to later exposure times could test if the population is a plateau (no significant difference) or a slow decline (significant difference). As we suspected, the long-term *E. coli* death curve slope is not zero, the cells are not in a plateau, and cells are dying slowly. This is evident by the fact that 3 h ampicillin treatment is significantly different (p<0.05) than the that longer exposures of ampicillin treatment: the p-values for 6 h, 8 h, and 24 h, were 6.5E-05, 4.6E-04, and 3.3E-06, respectively, compared to the 3 h (n≥3) (Table S4). There are points in the death curve where small plateaus (p>0.5) can be observed but overall, no plateau. This work underscores the importance of doing long-term kill curves as laid out in REF (Balaban et al., 2019). Next, we tested Syn3B P026 persisters and as expected, there was no plateau, the slope is not zero for different antibiotics (Fig. 2C), and there is a slow decline in persistence over time. This is evident by the fact that 24 h ciprofloxacin treatment was significantly different (p<0.05) than longer exposures of ciprofloxacin treatment: the p-values for 48 h and 72 h were 5.2E-05 and 3.6E-02 compared to the 24 h (n≥3). Similar results were obtained for streptomycin (100 µg/mL); the p-value for 48 h and 72 h treatment compared to 24 h is 4.2E-3 and 3.3E-2 (n≥3). Similar to *E. coli*, there were some points in the death curve that formed small plateaus (p>0.5), which again underscores the importance of doing long-term kill curves. Our results support a rapid decline in the short-term tolerance stage and then a slow decline in the persister stage, not a plateau, which aligns with the historical data (Bigger, 1944) and modern definition (Balaban et al., 2019).

After establishing that Syn3B forms persisters, we looked into genes previously shown to be related to tolerance in other organisms. We also find that the Syn3B genome lacks homologs of several genes and pathways reported in earlier research to be related to tolerance and persistence (Table S1). For instance, TA systems are often implicated for persistence and are still referenced as causing persistence (Moreno-del Álamo et al., 2020; Riffaud et al., 2020; Shenkutie et al., 2020), although recent research provided evidence of a lack of causation between persister formation and TA system (Conlon et al., 2016; Pontes and Groisman, 2019). In our study, we identify no known TA systems or homologous genes based on the results from TADB 2.0 (a database designed to search for TA systems based on homology) (Xie et al., 2018) in Syn3B. Our study clearly shows that TA systems are not required for tolerance or persistence in the minimal cell, and it seems likely that most bacteria do not require them either. It is not surprising that the minimal genome does not contain TA systems because the genome was designed by researchers and not subject to the natural environment. Natural TA systems are often described as “addictive” systems that are hard to lose once acquired; they often have overlapping toxin and antitoxin genes, making mutations less likely to be selected. Curiously, we observed some overlapping genes that are not at all similar to TA systems (they are also much longer in sequence than traditional TA systems, and are not homologous to any known TA system), and whose functions are not yet defined (Table S3).

Recently, the Wood group (Song and Wood, 2020; Wood and Song, 2020) proposed the ppGpp ribosome dimerization persister (PRDP) model for entering and exiting the persister state where ppGpp induces *hpf, rmf*, and *raiA*, which converts active ribosomes into their inactive state (such as 100s ribosome) to reduce translation, and consequently cells enter into persistence. Upon removal of antibiotic stress, cAMP level is reduced and HflX production is stimulated, which dissociates inactive 100S ribosomes into active 70S ribosomes and growth resumes. However, we did not detect homologs to the required genes, *raiA, rmf, hpf, and hflX*, for ribosome dimerization in the minimal cell genome, which means ribosome dimerization is not an essential mechanism for this organism (or another unknown mechanism for ribosome dimerization exists in Syn3B). (p)ppGpp may still play a role in Syn3B persistence, the genome contains *relA* (converts ATP and GTP to (p)ppGpp; Table 3). A potentially fruitful study would be to study the effects on survival by altering (p)ppGpp of Syn3B cultures.

This strain shows that TA systems and ribosome hibernation genes are not required for bacterial tolerance or persistence. Moreover, there are far fewer genes (less than 20 genes) present in Syn3B, which have been shown related to persistence. Thus, if specific genes or regulons are responsible for persistence, then Syn3B should be very useful in identifying them in future studies because it has a minimal number of genes. It is possible that persister formation and maintenance results from slowed or disrupted cellular networks (this is the hypothesis we most prescribe to), rather than the activity of specific genes or regulons. If tolerance and persistence results from slowed or disrupted cellular networks, then Syn3B is an ideal model organism to use because it has a minimal number of networks. While different genes and networks are likely responsible for persistence depending on the antibiotic, strain, or bacterial species, identifying genes in this minimal system should be applicable to a number of other microorganisms including the pathogen *Mycoplasma mycoides* from which Syn3B was derived. Regardless of how cells enter the dormant state, Syn3B provides a new model to study the genes and networks that allow cells to survive antibiotic treatment and could pave the way for finding new drugs that target the persister and tolerant subpopulations.

### Limitations of the study

We observed that despite controlling methodology to the greatest extent possible, persister levels varied considerably (far more than observed in our previous work with *E. coli*) (Deter et al., 2019a) between experiments for both streptomycin and ciprofloxacin treatments (Table S5). We hypothesize that this variability might be due to the stochastic fluctuations (noise) in gene expression levels, which results in protein level variations even among genetically identical cells in a similar environment (Soltani et al., 2016). We expect that gene expression and protein production to be more erratic in Syn3B compared to natural microorganisms because this cell has a designed genome lacking many control mechanisms and did not evolve to achieve some level of internal cellular “equilibrium” like native cells have.

We did not test for other subpopulations that have been identified during antibiotic treatment. We did not test for Syn3B transient tolerant cells, VBNCs, or spontaneous persisters.

## Methods

### Microbial strains and media

#### Mycoplasma mycoides

JCVI-Syn3B (Hutchison et al., 2016) and its derivatives were used in this study. Syn3B was a gift from Dr. John I. Glass from J. Craig Venter Institute, La Jolla, CA, USA. For evolution and antibiotic survival assays, cells were cultured at 37°C in SP16 media (57.5% 2X P1, 10.0% P4, 17.0% FBS, tetracycline 0.4% and vitamin B1 0.5%) (see Table S6 for details), which was developed based on SP4 media (Tully et al., 1979). All cultures were plated in SP4 agarose media (0.55% agarose) for colony counts. Note that agar is not used because it inhibits growth.

### Evolution by serial passage

Syn3B cultures were grown in 3 mL tubes at 37°C overnight in SP16 media. Overnight exponential cultures were serially passaged after each cycle of growth by transferring 30 µL of culture into 3 mL fresh media to initiate the next cycle of growth, and the cycles continued until a satisfactory growth rate was observed (Fig. S3. A). The culture was plated after 26 passages and a single colony was isolated, P026.

### Genome extraction, whole-genome sequencing, and identification of mutations

For whole-genome sequencing (WGS), a single colony of the evolved strains Syn3B P026 and all the resistant mutants (PK07_L1, PK07_L2, PS04_L1, PS04_L2, PC06_L1, PC06_L2, PSC09_L1, PSC09_L2) were isolated and inoculated into SP16 media for 24 h at 37°C. Next, genomic DNA was harvested and purified using Genomic DNA Purification Kit (ThermoFisher) in accordance with the manufacturer’s instructions. For quality checks, DNA purity and concentration were assessed by gel electrophoresis and Qubit Fluorimeter prior to sending for sequencing. Novogene Ltd. sequenced the genomes using paired-end Illumina sequencing at 2 × 150 bp read length and 350 bp insert size. A total amount of ∼1 μg of DNA per sample was used as input material for the DNA sample preparation. Sequencing libraries were generated from the qualified DNA samples using the NEB Next Ultra DNA Library Prep Kit for Illumina (NEB, USA) following the manufacturer’s protocol. For data processing, the original sequence data were transformed into raw sequenced reads by CASAVA base calling and stored in FASTQ (fq) format. For subsequent analysis, the raw data were filtered off the reads containing adapter and low-quality reads to obtain clean data. The resequencing analysis was based on reads mapping to reference genome of Syn3B by BWA software (Li and Durbin, 2009). SAMTOOLS (Li et al., 2009) was used to detect single nucleotide polymorphism (SNP) and InDels.

We also used Oxford Nanopore Technologies (ONT) MinION long-read sequencer to search for large insertions or gene duplication in all the evolved strains. ONT libraries were prepared using *Ligation kit* (SQK-LSK109) according to the manufacturer’s instructions. R 9.5 flow cell (FLO-MIN107, ONT) was used for sequencing based on manufacture protocol. The flow cell was mounted on a MinION Mk 1B device (ONT) for sequencing with the MinKNOW versions 19.12.5_Sequencing_Run_FLO-MIN107_SQK-LSK109 script. Then, reads were mapped against reference genome Syn3B using Geneious Prime software version 2020.1. (*https://www.geneious.com*). No large insertions were found in the sequenced genomes.

### Minimum inhibitory concentration (MIC) tests

Overnight cultures were serially diluted and plated onto SP4 agarose plates containing different concentrations of antibiotics (Strep, Cip, Cm, and Ksg) to determine the MIC of each antibiotic. Plates were incubated 4-5 days at 37°C before colony counts. MIC values were defined in this study as the lowest antibiotic concentration that inhibit the growth of Syn3B (See Fig. S4).

### Antibiotic survival assays

The schematic of the antibiotic survival assay from a stationary phase culture is shown in Fig. S3. C. Briefly, an overnight culture was diluted 1:100 into pre-warmed media and grown to exponential phase (OD_600_0.1-0.3). Next, the culture was separated into three flasks (for three biological replicates) and grown at 37°C until it reached stationary phase (OD 0.45-0.55, which takes ∼3-6 h). After that, each culture was diluted 1:10 into 100 µg/mL streptomycin (10X MIC) or 1 µg/mL (10X MIC) ciprofloxacin containing pre-warmed media and kept at 37°C shaking at 250 rpm for 72 h. Samples were taken at different time points until 72 h for the time-kill assays. To remove the antibiotic before plating, 100 μl of each sample was washed with 1.9 mL ice-cold SP16 media and collected by centrifugation (16,000 rpm for 3 min at 4°C). Cells were then resuspended and serially diluted into ice-cold SP16 media and plated to count the colony-forming units (CFU). Persisters were quantified by comparing CFUs per milliliter (CFU/ml) before antibiotic treatment to CFU/ml after antibiotic treatment. Plates were incubated at 37°C for 4-5 days, then scanned using a flatbed scanner (Datla et al., 2017; Deter et al., 2019a; Levin-Reisman et al., 2017). Custom scripts were used to identify and count bacterial colonies (Deter et al., 2019a; Deter et al., 2019b) used to calculate CFU/ml and persister frequency. Over 200 colonies were streaked periodically into antibiotic-containing plates to test for antibiotic-resistant mutants. Also, antibiotic-treated culture was washed after 48 h, regrew to stationary phase, and exposed the culture again in the same antibiotic treatment for 48 h and plated after 24 h and 48 h to observe the difference between the first (Strep/Cip) and second death curve (Strep re-exposed/Cip re-exposed) in both antibiotic treatment. Persister assay for *E*.*coli* 24 h ampicillin treatment (unpublished data) was done by similar manner as described in REF (Deter et al., 2020a)

### Evolution through cyclic antibiotic treatment

Stationary phase culture was exposed to 100 μg/mL streptomycin (∼10× MIC), 1 μg/mL ciprofloxacin (∼10× MIC), a combination of streptomycin (100 μg/mL) and ciprofloxacin (1 μg/mL) and 300 μg/mL kasugamycin (∼10× MIC) antibiotic for 24 h, then antibiotic-containing medium was removed by washing twice with SP16 medium (10 min of centrifugation at 7000 g at 4°C). Finally, the culture was resuspended in 10 mL of fresh SP16 media and grown overnight at 37 °C. After every cycle of antibiotic treatment, the tolerance phenotype was observed. Finally, we isolated single colony from evolved populations from streptomycin (PS04), ciprofloxacin (PC06), combination of streptomycin and ciprofloxacin (PSC09) and kasugamycin (PK07) treatment after four, six, nine and seven cycle, respectively, and then their genomes were sequenced. Two different evolutionary lineages were used for all evolved populations.

### Determination of growth and doubling times

Overnight cultures were diluted into OD 0.1 (measured in SpectronicTM 200E) and 30 μL of diluted cultures were inoculated into individual wells containing 270 µL of SP16 media in a 96-Well Optical-Bottom Plate with Polymer Base (ThermoFisher) to measure OD at 600 nm using FLUOstar Omega microplate reader. Doubling time was determined by the linear regression of the natural logarithm of the OD over time during exponential growth as described in REF (Widdel, 2007).

### Statistical analysis

All data is presented in the manuscript as mean ± SEM of at least three independent biological replicates. Statistical significance was assessed using an f-test to determine variance (p < 0.05 was considered to have significant variance), followed by a two-tailed t-test with unequal variances (if F statistic > F critical value) or equal variances (if F statistic < F critical value).

## Acknowledgments

*Mycoplasma mycoides* JCVI-Syn3B was a generous gift from Dr. John I. Glass from J. Craig Venter Institute (JCVI), La Jolla, CA, USA. This work is supported by the National Science Foundation award Numbers 1922542 and 1849206, and by a USDA National Institute of Food and Agriculture Hatch project grant number SD00H653-18/project accession no. 1015687.

## Author contributions

T.H. wrote the manuscript and performed most experiments. H.S.D. repeated streptomycin persister assay and developed custom code for colony counting, growth rate, and statistical analyses. E.P. performed cyclic antibiotic treatment experiments. N.C.B. planned and directed the project. All authors contributed to discussing and editing the manuscript.

## Data and materials availability

All data that supports the findings of this study are available from the corresponding author upon request. Whole genomes data of P026, PK07_L1, PK07_L2, PS04_L1, PS04_L2, PC06_L1, PC06_L2, PSC09_L1 and PSC09_L2 strains has been deposited on the NCBI Genome Bank in BioProject PRJNA635211 with the accession number CP053944, CP069339, CP069340, CP069341, CP069342, CP069343, CP069344, CP069345, CP069346, respectively. The code used for colony counting is available on GitHub at *https://github.com/hdeter/CountColonies*.

## Supplementary materials

**Fig. S1.**
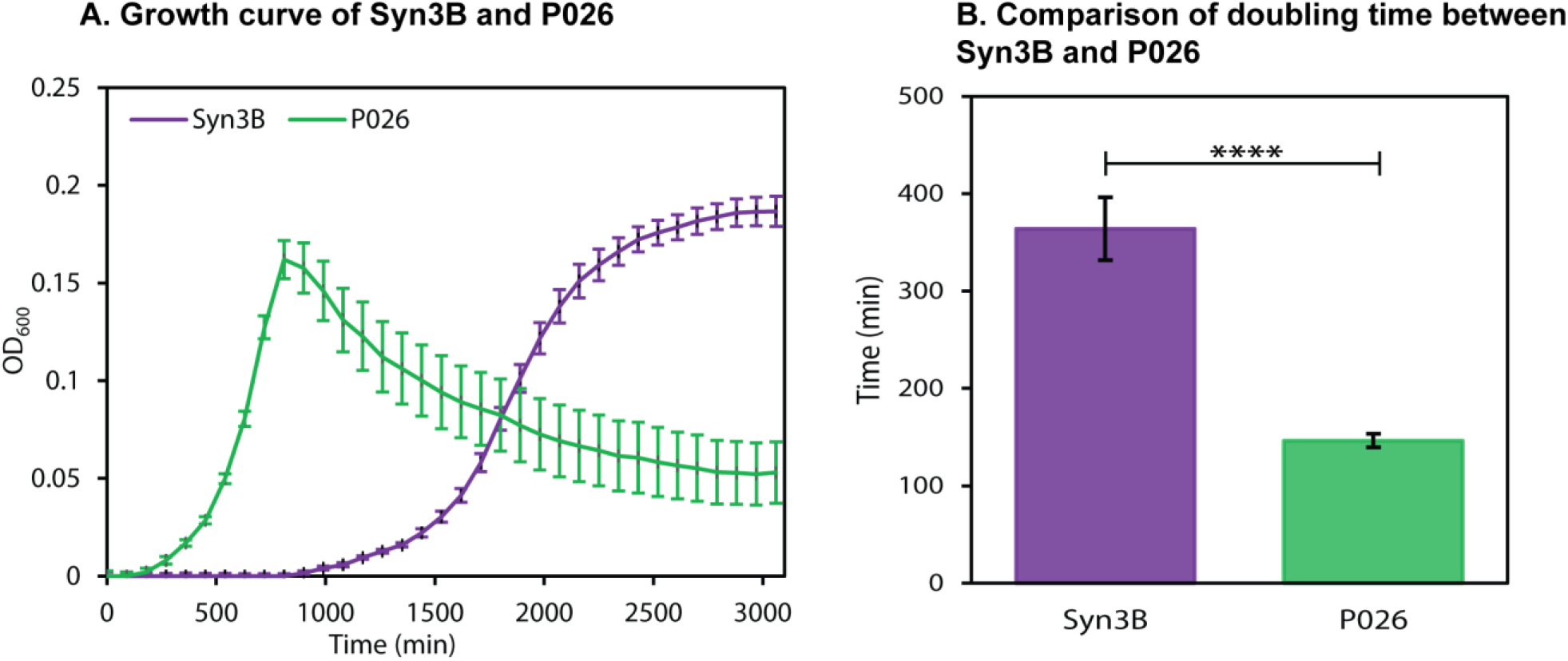
Growth of the minimal cell. **(A)** Growth curve and **(B)** doubling time of evolved strain Syn3B P026 and parent strain Syn3B. Error bar represents SEM. n = 12 independent biological replicates. *p<0.05, **p<0.01, ***p<0.001, ****p < 0.0001.

**Fig. S2.**
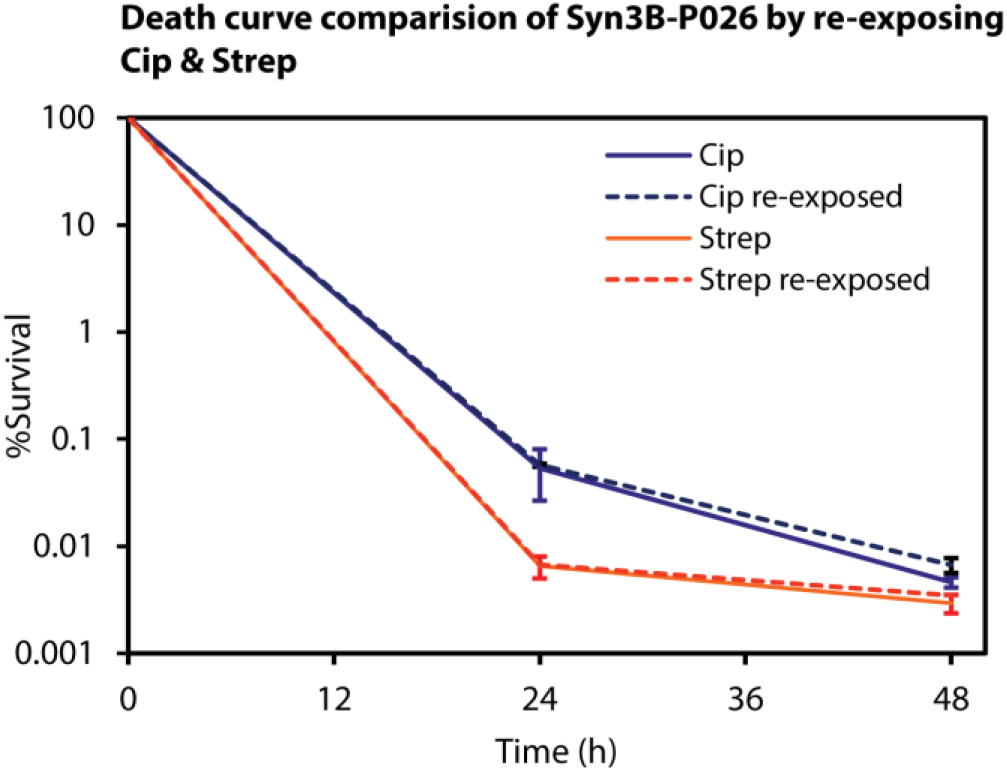
Death curve comparison of Syn3B P026 by re-exposing streptomycin and ciprofloxacin. Overnight cultures of Syn3B were grown to stationary phase (OD∼0.3-0.35), diluted to 1:10 and treated with streptomycin (100 µg/mL) or ciprofloxacin (1 µg/mL) for 48 h and sampled after 24 h and 48 h. Antibiotic treated culture then washed twice through centrifugation, grew back to stationary phase and re-exposed to same antibiotic to make the second death curve. Error bars represent SEM (n ≥ 3).

**Fig. S3.**
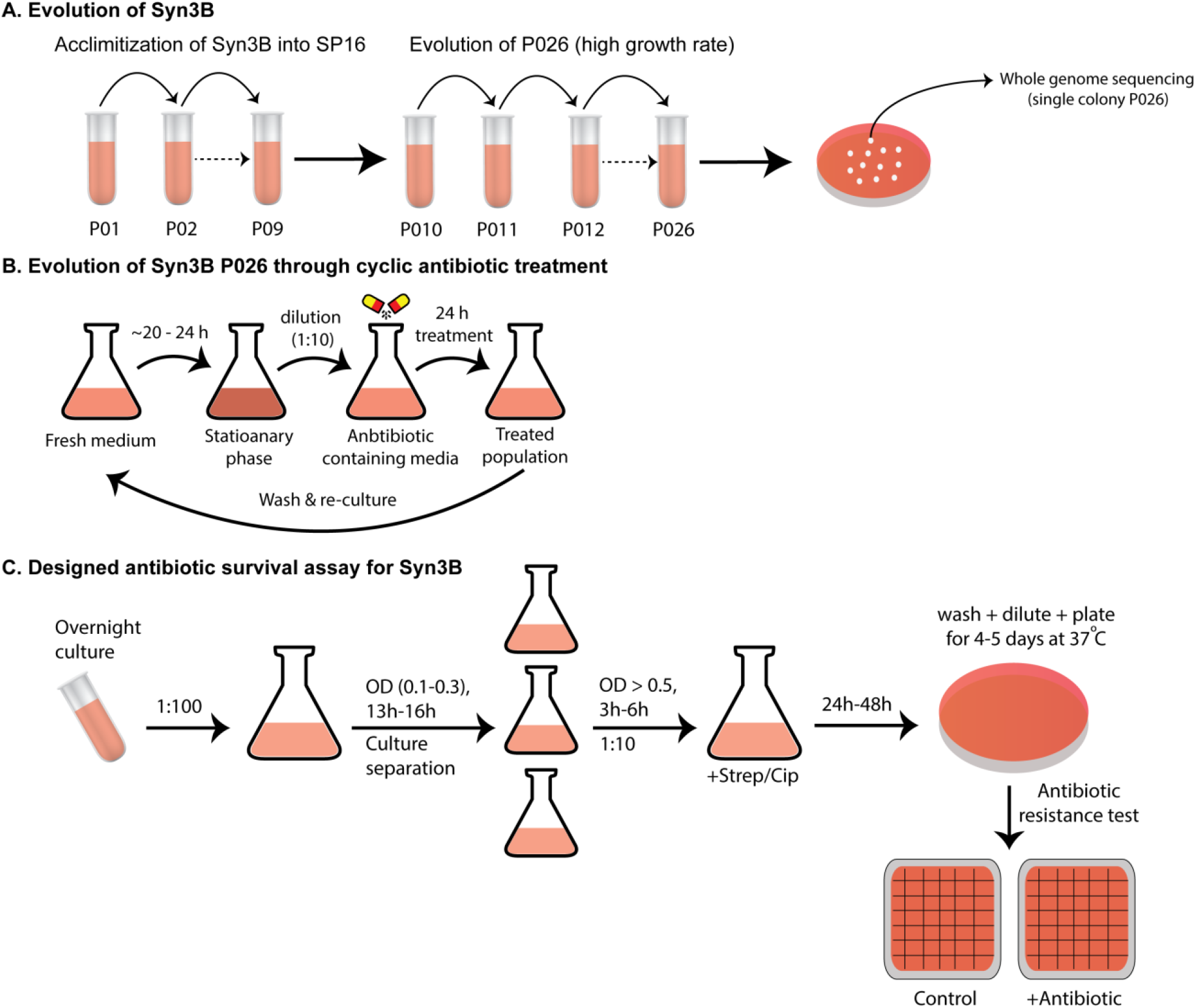
**A**. Evolution of Syn3B P026. Cells were evolved in SP16 media by a serial passage from P01 to P09. From P09, exponential cultures were repeatedly diluted 1:100 into fresh SP16 media for higher growth yield up until the 26 passage. Then, a single colony called P026 was selected and send for sequencing. **B**. Evolution of Syn3B resistant mutants through cyclic antibiotic treatment. Stationary phase cultures of P026 were diluted 1:10 in SP16 media containing lethal doses of different types of antibiotic (Streptomycin, Ciprofloxacin, Streptomycin-Ciprofloxacin and Kasugamycin) for 24 h, then washed twice through centrifugation, regrew in the similar condition and re-exposed with same antibiotic. Finally, a single colony was selected for whole genome sequencing. **C**. Antibiotic survival assay was optimized based on traditional agar plate method. Overnight cultures were grown to stationary phase (OD>0.5), diluted to 1:10 in antibiotic-containing media. Percent surviving cells were calculated by the counting colony number before and after antibiotic treatment. Over 200 individual colonies were tested for bacterial resistance, and as expected, no resistant colonies were detected.

**Fig. S4.**
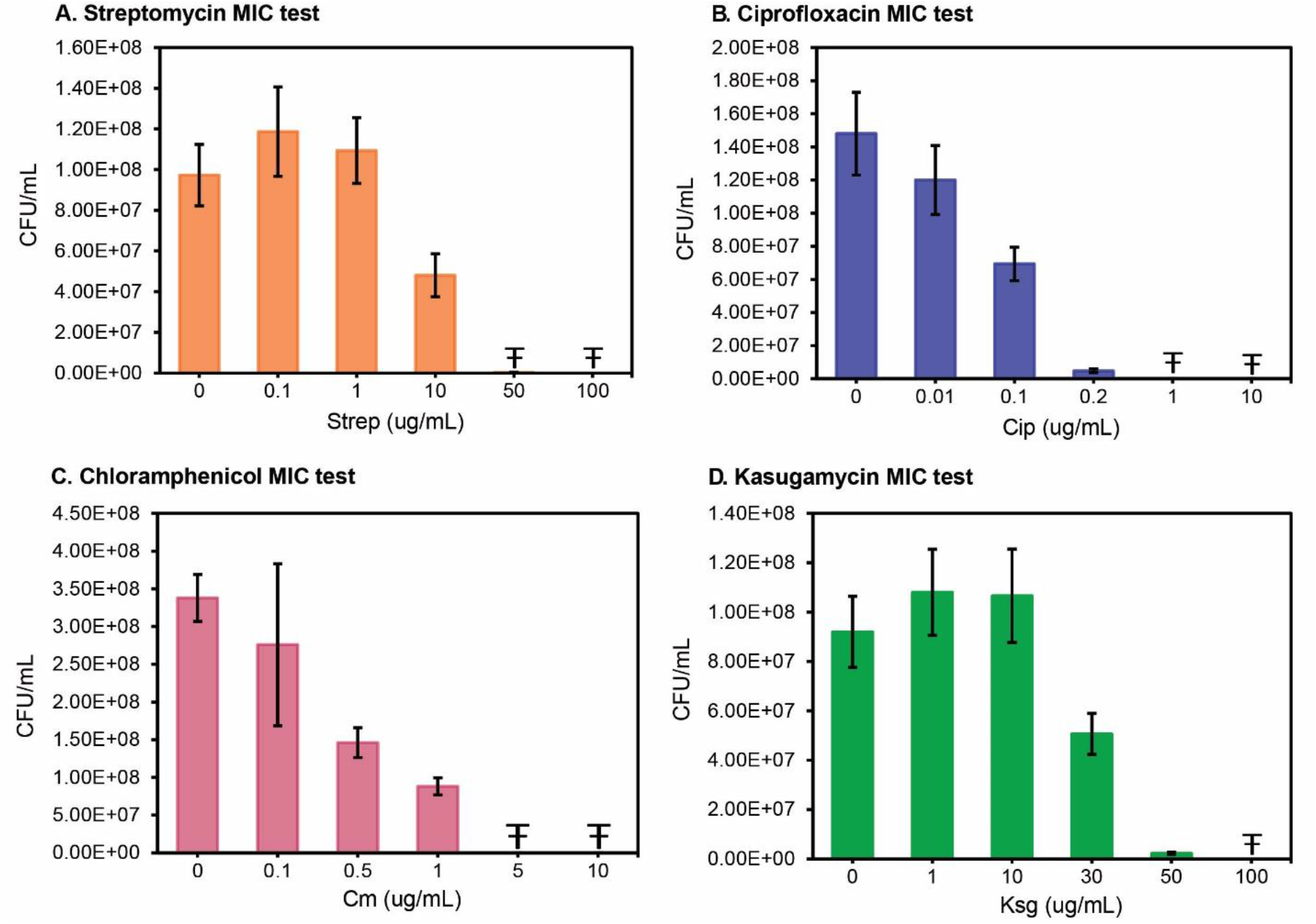
Determination of Minimal Inhibitory Concentration (MIC). **A**. Streptomycin (Strep) **B**. Ciprofloxacin (Cip). **C**. Chloramphenicol (Cm). **D**. Kasugamycin (Ksg). Exponential phase cultures with different dilutions were plated on SP4 agarose plate with different concentrations of antibiotics. The MIC was determined to be 10 μg/ml for streptomycin, 0.1 µg/mL for ciprofloxacin, 0.5 µg/mL for chloramphenicol, and 30 µg/mL for kasugamycin. Error bars represent the standard deviation and Ŧ represents data is out of detectable range.

**Table S1.**
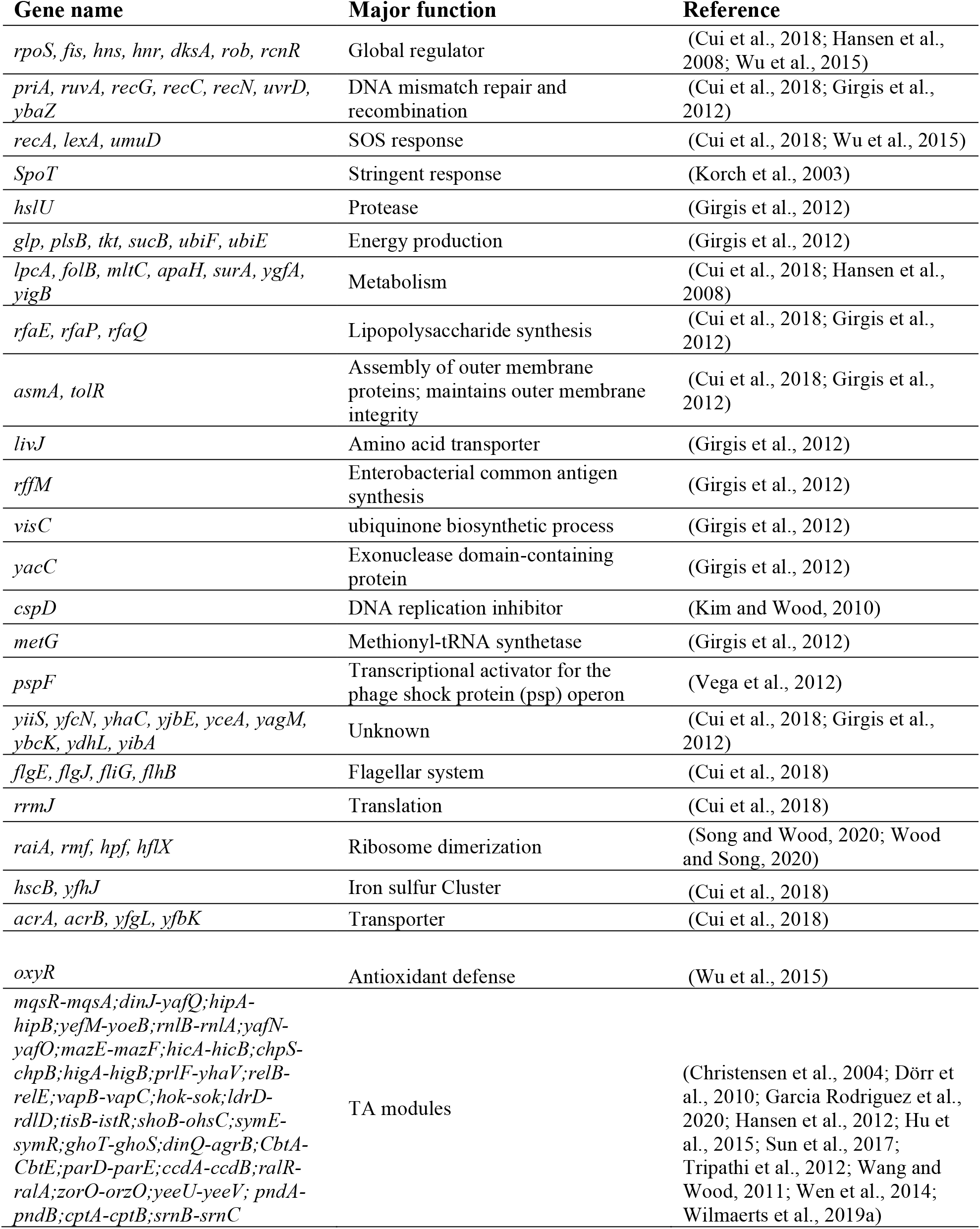
**The** Syn3B genome is lacking homologs to stress response genes that are linked with tolerance and persistence.

**Table S2.** Comparison of % similarity (based on homologous proteins) of Syn3B with most threatening human pathogens.

| Human pathogens | Related diseases | Homologous proteins No. in Syn3B* | % similarity with Syn3B |
| --- | --- | --- | --- |
| *Escherichia coli* O157 | Severe intestinal infection | 287 | 63 |
| *Staphylococcus aureus* | Pneumonia, meningitis, osteomyelitis, endocarditis, toxic shock syndrome, bacteremia, sepsis, impetigo, boils, cellulitis, folliculitis and carbuncles etc. | 338 | 74 |
| *Burkholderia cepacia* | pneumonia in immunocompromised individuals | 262 | 58 |
| *Pseudomonas aeruginosa* | Urinary tract infections, respiratory system infections and dermatitis | 273 | 60 |
| *Clostridium difficile* | Diarrhea and inflammation of the colon | 329 | 73 |
| *Klebsiella pneumoniae* | Pneumonia and infection in the lungs | 294 | 65 |
| *Acinetobacter baumannii* | Infection in the blood, urinary tract, and lungs (pneumonia) | 272 | 60 |
| *Mycobacterium tuberculosis* (MTB) | Tuberculosis | 275 | 60 |
| *Neisseria gonorrhoeae* | Gonorrhea | 265 | 58 |
| *Streptococcus pyogenes* | Pharyngitis, tonsillitis, scarlet fever, cellulitis, erysipelas and rheumatic fever etc. | 328 | 72 |

**Table S3.** Overlapping genes in Syn3B.

| Gene name | Gene Length (bp) | Locus tag | Annotation |
| --- | --- | --- | --- |
| *ietA* | 1062 | JCVISYN3A_0132 | AAA family ATPase |
| *ietS* | 2,274 | JCVISYN3A_0133 | hypothetical protein |
| *CDS_33* | 681 | JCVISYN3A_0281 | hypothetical protein |
| *proRS* | 1425 | JCVISYN3A_0282 | Proline--tRNA ligase |
| *truB* | 879 | JCVSYN2_00715 | tRNA pseudouridine(55) synthase TruB |
| *ribF* | 555 | JCVSYN2_00720 | FAD synthetase |
| *CDS_50* | 741 | JCVSYN2_00955 | hypothetical protein |
| *CDS_51* | 345 | JCVSYN2_00960 | hypothetical protein |
| *trmK*; *yqfN* | 678 | JCVSYN2_01080 | hypothetical protein |
| *CDS_60* | 777 | JCVSYN2_01085 | dinuclear metal center protein, YbgI family |
| *pncB* | 1062 | JCVSYN2_01655 | Nicotinate phosphoribosyltransferase |
| *CDS_97* | 609 | JCVSYN2_01660 | Uncharacterized protein |
| *CDS_124* | 681 | JCVSYN2_02430 | hypothetical protein |
| *CDS_125* | 1539 | JCVSYN2_02435 | amino acid permease |
| *CDS_99* | 363 | JCVSYN2_01695 | lipoprotein |
| *CDS_100* | 1071 | JCVSYN2_01700 | hypothetical protein |

**Table S4.**
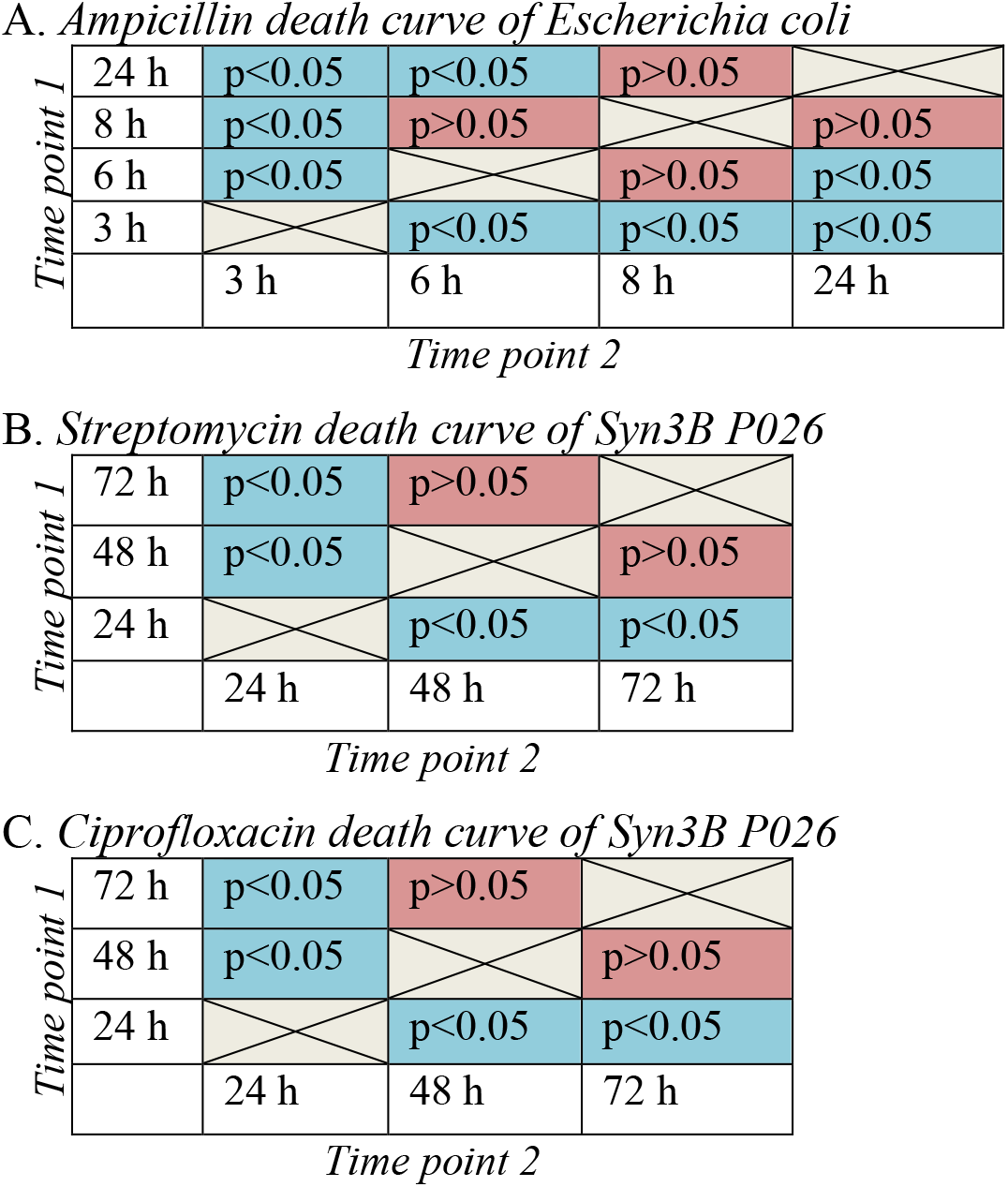
Statistical analysis between different time points of ampicillin death curve for *Escherichia coli* (A), streptomycin death curve for Syn3B P026 (B) and ciprofloxacin death curve for Syn3B P026 (C). Statistical significance was assessed using an f-test to determine variance (p < 0.05 was considered to have significant variance), followed by a two-tailed t-test with unequal variances (if F statistic > F critical value) or equal variances (if F statistic < F critical value). A. *Ampicillin death curve of Escherichia coli*

**Table S5.**
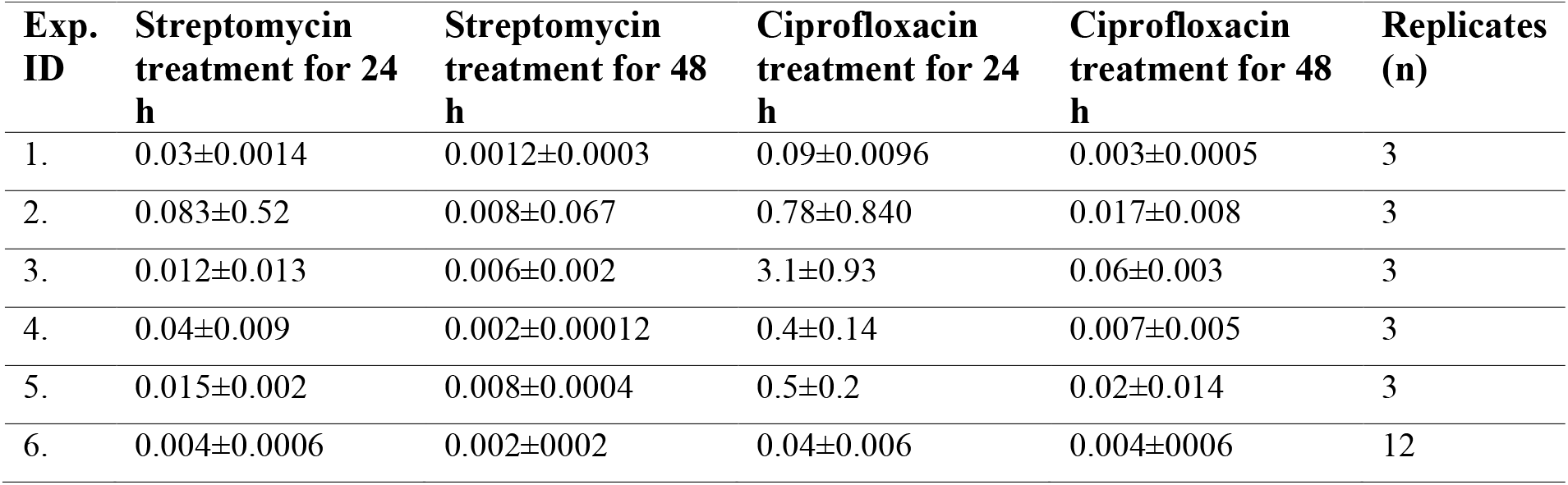
Experimental variation in population decay curve of Syn3B P026.

**Table S6.** SP16 media composition.

| **SP16 media components** |  |  |  |
| --- | --- | --- | --- |
| **Components** | **Final** | **Supplier** | **Cat No.** |
| **2XP1** |  |  |  |
| Mycoplasma Broth Base (PPLO Broth) | 7 mg/ml | Thomas Scientific | BD 211458 |
| Tryptone | 20 mg/ml | Fisher | AC611841000 |
| Peptone | 10.6 mg/ml | Fisher | BP9725-500 |
| Yeast extract solution (autoclaved) | 14 mg/ml | Fisher | BP9727-500 |
| TC Yeastolate (Autoclaved) | 4 mg/ml | Thomas Scientific | BD 255772 |
| **P4** |  |  |  |
| D(+)-Glucose | 0.50% | Fisher | Alfa Aesar A1682836 |
| CMRL 1066 (with L-Glutamine, without Sodium bicarbonate) | 0.25X | Thomas Scientific | C992B09 Mfr. No. AT110-1L |
| NaHCO<sub>3</sub> (Sodium Bicarbonate) | 1.1 mg/ml | Fisher | S233-3 |
| Penicillin G | 625 µg/ml (~1000 U/ml) | Fisher | MP Biomedicals 0210054380 (powder: 500-1700 u/mg) |
| L-Glutamine | 146 µg/ml | Fisher | Alfa Aesar A1420118 |
| **FBS** |  |  |  |
| Fetal Bovine Serum (FBS), Heat inactivated | 17% | Fisher | 10-438-018 Gibco 10438018 |
| Tetracycline | 4 µg/ml | Fisher | BP912-100 |
| Vit B1 | 5 µg/ml | Acros organic | 148990100 |

